# Aberrations in Notch-Hedgehog signalling reveal cancer stem cells harbouring conserved oncogenic properties associated with hypoxia and immunoevasion

**DOI:** 10.1101/526202

**Authors:** Wai Hoong Chang, Alvina G. Lai

## Abstract

**Background:** Cancer stem cells (CSCs) have innate abilities to resist even the harshest of therapies. To eradicate CSCs, parallels can be drawn from signalling modules that orchestrate pluripotency. Notch-Hedgehog hyperactivation are seen in CSCs, yet, not much is known about their conserved roles in tumour progression across cancers.

**Methods:** Employing a comparative approach involving 21 cancers, we uncovered clinically-relevant, pan-cancer drivers of Notch and Hedgehog. GISTIC datasets were used to evaluate copy number alterations. Receiver operating characteristic and Cox regression were employed for survival analyses.

**Results:** We identified a Notch-Hedgehog signature of 13 genes exhibiting high frequencies of somatic amplifications leading to transcript overexpression. The signature successfully predicted patients at risk of death in five cancers(n=2,278): glioma(P<0.0001), clear cell renal cell(P=0.0022), papillary renal cell(P=0.00099), liver(P=0.014) and stomach(P=0.011). The signature was independent of other clinicopathological parameters and offered additional resolution to stratify similarly-staged tumours. High-risk patients exhibited features of stemness and had more hypoxic tumours, suggesting that hypoxia may influence CSC behaviour. Notch-Hedgehog^+^ CSCs had an immune privileged phenotype associated with increased regulatory T cell function.

**Conclusion:** This study will set the stage for exploring adjuvant therapy targeting the Notch-Hedgehog axis to help optimise therapeutic regimes leading to successful CSC elimination.

## Background

Tumours are far from homogeneous masses, yet many contemporary therapies continue to treat them as such. It has become increasingly clear that a minor population of tumour cells known as cancer stem cells (CSCs) contribute to treatment resistance as they have the propensity to tolerate DNA damage^1,2^ and evade immune detection^3^ to give rise to new tumours post therapy. Identification of CSCs has remained a challenging endeavour since they only make up a small proportion of the tumour and are histologically similar to non-stem cancer cells. Moreover, molecular markers that identify CSCs are often cancer-type dependent, which limit their broad scale applications^4^. CSCs share many qualities with embryonic or adult stem cells. For example, activation of signalling pathways involved in coordinating cellular homeostasis, morphogenesis and cell fate determination (TGF-β, Wnt, Notch and Hedgehog) are often seen in CSCs. These pathways rarely act in isolation and significant crosstalk between them have been reported^5^.

In order to fully exploit these pathways for CSC therapy, pan-cancer explorations are warranted to reveal conserved components that can be prioritised as therapeutic targets. Concentrating on Notch and Hedgehog signalling pathways, we seek to attain a comprehensive understanding of how somatic copy number alterations and expression profiles of pathway genes along with their downstream targets could influence tumour progression and prognosis. The role of Notch signalling in oncogenesis was initially discovered in T cell acute lymphoblastic leukaemia^6^. Since then, multiple studies on Notch signalling have demonstrated both oncogenic and tumour suppressive functions in haematological and solid malignancies, implying its pleiotropic nature that is very much dependent on cellular types^7^. Hedgehog is a morphogen that regulates a signalling cascade involving the Smoothened protein to influence morphogenetic processes such as proliferation and differentiation^8^. Interactions between Notch and Hedgehog signalling have been demonstrated in multiple cancers. *Hes1*, a Notch effector, is targeted by *sonic hedgehog* in neural cells^9^. When *Patched*, a negative regulator of Hedgehog, is abrogated in mice, this gives rise to medulloblastoma with enhanced Notch signalling^10^. Hedgehog signalling promotes the expression of *Jagged2 (*a Notch ligand)^11^ and in ovarian cancer mice models, inhibition of *Jagged1* would sensitise tumours to docetaxel treatment by affecting *GLI2* function^12^. Concurrent activation of Hedgehog and Notch signalling was observed in prostate cancer cells that were resistant to docetaxel^13^. Glioblastoma treated with a Notch inhibitor was subsequently desensitised to further Notch suppression as they upregulate Hedgehog signalling^14^.

These studies highlight the importance of Notch-Hedgehog interactions in cancer, which calls for a better understanding of their relationship and also to reveal crosstalk with other pathways involved in regulating CSC function. Harnessing genomic and transcriptomic sequences of 21 cancer types, we performed a comprehensive investigation linking genomic alterations to transcriptional dysregulation of Notch-Hedgehog pathway genes. We discovered conserved patterns of Notch-Hedgehog hyperactivation across cancers and revealed putative driver genes that were associated with CSC phenotypes underpinning poor clinical outcomes. We also examined the relationship between the tumour microenvironment (hypoxia and immune suppression) and Notch-Hedgehog^+^ CSCs. In-depth knowledge of the Notch-Hedgehog signalling axis afforded by this study will set the stage for exploring combinatorial chemotherapy targeting both pathways simultaneously to potentially eradicate CSCs.

## Materials and Methods

A total of 72 genes associated with Notch and Hedgehog signalling were retrieved from the Kyoto Encyclopedia of Genes and Genomes (KEGG) database listed in Table S1.

### Study cohorts

We retrieved transcriptomic and genomic profiles of 21 cancer types (n=18,484) including their non-tumour counterparts from The Cancer Genome Atlas and Broad Institute GDAC Firehose^15^ (Table S2).

### Somatic copy number alterations analyses

We retrieved Firehose Level 4 copy number variation datasets in the form of GISTIC gene-level tables, which provided discrete amplification and deletion indicators^16^. A sample was defined as ‘deep amplification’ for values that were higher than the maximum median copy-ratio for each chromosome arm (> +2). Samples with values less than the minimum median copy-ratio for each chromosome arm were called ‘deep deletions’ (< -2). GISTIC indicators of +1 and -1 represented shallow amplifications and deletions respectively.

### Calculating Notch-Hedgehog 13-gene scores, hypoxia scores and regulatory T cell (Treg) scores

The Notch-Hedgehog 13-gene signature was employed to calculate a score for each patient. It comprised of the following genes: *JAG1, LFNG, DTX2, DLL3, GPR161, PSENEN, GLI1, HES1, PTCRA, DTX3L, ADAM17, KIF7* and *NOTCH1*. Hypoxia scores were calculated from 52 hypoxia signature genes^17^. Treg scores were calculated based on the overlap between four Treg signatures^18^–^21^, consisting of 31 genes: *FOXP3, TNFRSF18, TNFRSF9, TIGIT, IKZF2, CTLA4, CCR8, TNFRSF4, IL2RA, BATF, IL2RB, CTSC, CD27, PTTG1, ICOS, CD7, TFRC, ERI1, GLRX, NCF4, PARK7, HTATIP2, FCRL3, CALM3, DPYSL2, CSF2RB, CSF1, IL1R2, VDR, ACP5* and *MAGEH1*. Scores were calculated from the average log_2_ expression values of 13, 52 or 31 genes representing Notch-Hedgehog, hypoxia and Tregs respectively. Kaplan-Meier analyses of the Notch-Hedgehog signature were performed on patients separated into quartiles based on their 13-gene scores. For analyses in Figures 4, 5 and 6, patients were separated into four groups using median 13-gene scores and median CSC transcription factor expression levels (*EZH2, REST* and *SUZ12*), hypoxia scores or Treg scores as thresholds for Kaplan-Meier and Cox regression analyses. Nonparametric Spearman’s rank-order correlation tests were used to investigate the relationship between 13-gene scores and TF expression levels, hypoxia scores or Treg scores.

### Multidimensional scaling, differential expression and survival analyses

As per the journal’s guidelines, we have not repeated methods here as we have previously published detail methods for multidimensional scaling (MDS), differential expression and survival analyses^22^. Briefly, MDS analysis was employed to visualise samples’ distance (tumour and non-tumour) in reduced 2-dimensional space. The R vegan package was employed for MDS ordination using Euclidean distances. Permutational multivariate analysis of variance (PERMANOVA) was used to investigate statistical differences between tumour and non-tumour samples. The linear model and Bayes method was employed for differential expression analyses, followed by the Benjamini-Hochberg false discovery rate method. Kaplan-Meier, Cox proportional hazards and receiver operating characteristic survival analyses were performed using R survminer, survival and survcomp packages.

### Functional enrichment and transcription factor (TF) analyses

Differential expression analyses as mentioned previously were performed on patients separated into quartiles 4 and 1 based on their 13-gene scores. Differentially expressed genes were mapped against KEGG and Gene Ontology (GO) databases using GeneCodis^23^ to determine pathways that were enriched. The Enrichr tool was used to determine whether differentially expressed genes were enriched for stem cell TFs binding targets by comparing chromatin immunoprecipitation sequencing profiles from ChEA and ENCODE databases^24^.

The R ggplot2 and pheatmap packages were used to generate all plots.

## Results

### Recurrently amplified driver genes associated with Notch and Hedgehog activation in 21 diverse cancer types

To characterise the extent of Notch and Hedgehog signalling and identify common molecular subtypes, we examined somatic copy number alterations (SCNAs) and differential expression (tumour versus non-tumour) patterns of 72 genes in 18,484 cases of clinically annotated stage I to IV samples representing 21 cancer types (Fig. 1A; Table S1; Table S2). We found that 70 out of 72 genes were recurrently amplified in at least 20% of samples per cancer type in at least one cancer type (Fig. 1A). Lung squamous cell carcinoma (LUSC) had the highest fraction of samples harbouring amplified Hedgehog genes, while endometrial cancer (UCEC) had the fewest somatic gains (Fig. 1B). When considering Notch gene amplifications, LUSC also emerged as the top candidate while clear cell renal cell carcinoma (KIRC) had the fewest number of Notch gene amplifications (Fig. 1B). In terms of focal deletions, this was also the highest in LUSC for Hedgehog genes and renal chromophobe carcinoma (KICH) for Notch genes (Fig. 1B).

**Figure 1.**
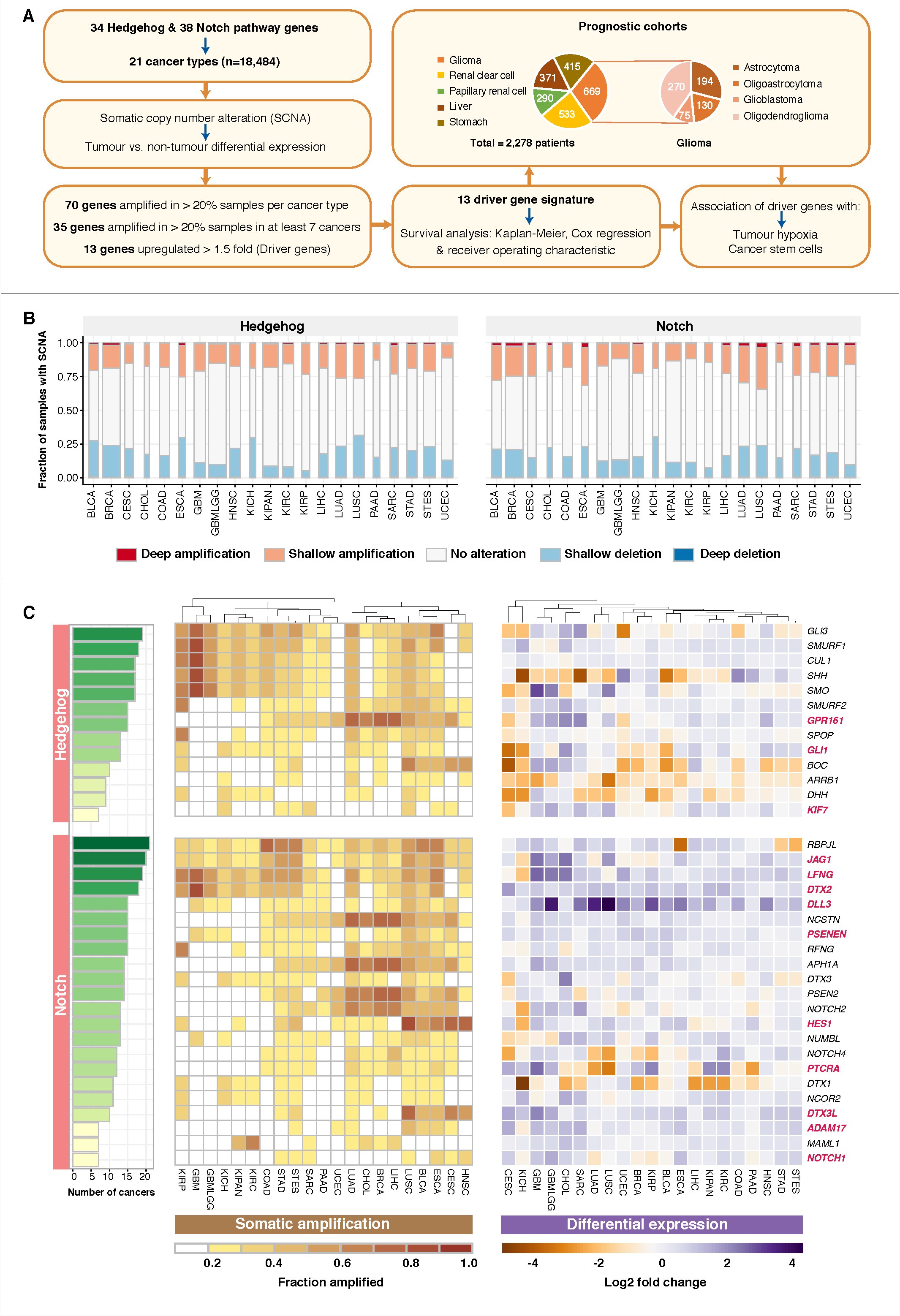
Pan-cancer drivers of Notch-Hedgehog signalling. **(A)** Schematic diagram illustrating the study design and the identification of Notch-Hedgehog driver genes, which represent the 13-gene signature. SCNA and expression profiles of 72 Notch-Hedgehog pathway genes were interrogated in 21 cancer types involving 18,484 patients. We identified 70 genes as amplified in at least 20% of samples and 35 genes that were amplified in at least 20% of samples in at least 7 cancer types. Differential expression analyses between tumour and non-tumour samples revealed that the 13 recurrently amplified genes were also upregulated, potentially indicating a gain-of-function. These 13 genes were prioritised as a Notch-Hedgehog signature, which was prognostic in five cancer types involving 2,278 patients. Associations of the signature with tumour microenvironmental features of hypoxia and immunity were also investigated. Pie slices indicate the number of samples within each cancer type. (**B)** Stacked bar graphs represent the proportion of samples in each cancer type with SCNA of Hedgehog and Notch pathway genes. Width of the bars reflect the number of samples within each cancer. (**C)** Somatic gains and differential expression profiles of 35 Notch-Hedgehog genes that were recurrently amplified in at least 7 cancer types (one-third of all cancers). Cumulative bar charts on the left represent the number of cancers with at least 20% of samples with somatic amplification. Heatmap on the left represents the extent of somatic gains for each of the 35 genes separated into Hedgehog and Notch signalling pathways across 21 cancers. Heatmap intensities depict the fraction of the cohort in which a given gene is amplified. Columns (cancer types) were ordered using Euclidean distance metric and hierarchical clustering to reveal cancers that were similar. Heatmap on the right represents tumour and non-tumour differential expression values (log_2_) for the 35 genes. Genes highlighted in red represent the 13 Notch-Hedgehog signature genes. Cancer abbreviations were listed in Table S2.

Focusing on recurrently amplified genes, we identified 35 genes (Hedgehog pathway: 13 genes; Notch pathway: 22 genes) that were gained in >20% of samples and in at least one-third of cancer types (> 7 cancers) (Fig. 1C). *GLI3, SMURF1, RBPJL, JAG1, LFNG* and *DTX2* were some of the most amplified genes present in at least 18 cancers (Fig. 1C). In contrast, *KIF7, NOTCH1, MAML* and *ADAM17* were the least amplified genes (Fig. 1C). LUSC had the highest number of amplified genes (34 genes) followed by 33 genes in oesophageal carcinoma (ESCA) and stomach and oesophageal carcinoma (STES) and 32 genes in stomach adenocarcinoma (STAD) and bladder urothelial carcinoma (BLCA) (Fig. 1C). In contrast, only 8 genes were amplified in UCEC (Fig. 1C).

SCNA events associated with overexpression could represent candidate driver genes since positive correlations between gene amplification and overexpression are indicative of a gain-of-function^25^. Differential expression analyses between tumour and adjacent non-tumour samples revealed that 13 of the amplified genes were also significantly upregulated (> 1.5 fold-change, P<0.05) in tumours of at least 7 cancer types (Fig. 1C). These genes were prioritised as a Notch-Hedgehog signature potentially representative of multiple cancers: *JAG1, LFNG, DTX2, DLL3, GPR161, PSENEN, GLI1, HES1, PTCRA, DTX3L, ADAM17, KIF7* and *NOTCH1 (*Fig. 1C).

### A 13-gene Notch-Hedgehog signature predicts survival outcomes in five cancers

Tumours displayed various degrees of somatic gains and overexpression of Notch-Hedgehog pathway genes (Fig. 1), suggesting that aberrant activation of these pathways may influence disease progression and survival outcomes. We employed univariate Cox proportional hazards regression analyses to test the prognostic roles of individual Notch-Hedgehog signature genes across 20 cancer types where survival data is available. Prognosis appeared to tissue type-dependent (Fig. S1). All 13 genes were prognostic in the glioma dataset (GBMLGG), consisting of samples from patients with astrocytoma, oligoastrocytoma, oligodendroglioma and glioblastoma multiforme (Fig. S1). A majority of the genes (9 out of 13) were associated with poor prognosis (hazard ratio [HR] > 1, P<0.05) (Fig. S1). However, despite showing high frequencies of SCNAs (Fig. 1C), none of the 13 genes harboured prognostic information in patients with LUSC, cholangiocarcinoma (CHOL) or oesophageal carcinoma (ESCA) (Fig. S1).

We next considered all 13 genes as a group in assessing prognosis. For each patient, we calculated their 13-gene scores by taking the average expression of all genes. Patients were separated into survival quartiles based on their 13-gene scores. Remarkably, Kaplan-Meier estimates and log-rank tests revealed that the 13-gene signature accurately predicted patients at higher risk of death in five cancer types (n=2,278): glioma (P<0.0001), clear cell renal cell (P=0.0022), papillary renal cell (P=0.00099), liver (P=0.014) and stomach (P=0.011) (Fig. 2A). Patients within the 4^th^ quartile had significantly poorer survival rates compared to those within the 1^st^ quartile: glioma (HR=3.386, P<0.0001), clear cell renal cell (HR=2.177, P=0.00048), papillary renal cell (HR=4.881, P=0.0053), liver (HR=2.627, P=0.0039) and stomach (HR=2.217, P=0.014) (Table S3). When comparing tumour and non-tumour expression patterns, Mann-Whitney-Wilcoxon tests revealed that a vast majority of the 13 genes were significantly upregulated in tumours of these cancers (Fig. S2) where hyperactivation of Notch-Hedgehog signalling was associated with adverse survival outcomes (Fig. 2A). Multidimensional scaling analyses revealed that the 13 genes could accurately distinguish tumour from non-tumour samples in these cancers (Fig. 2B), suggesting that Notch-Hedgehog transcriptional states could be used to identify cells with oncogenic properties.

**Figure 2.**
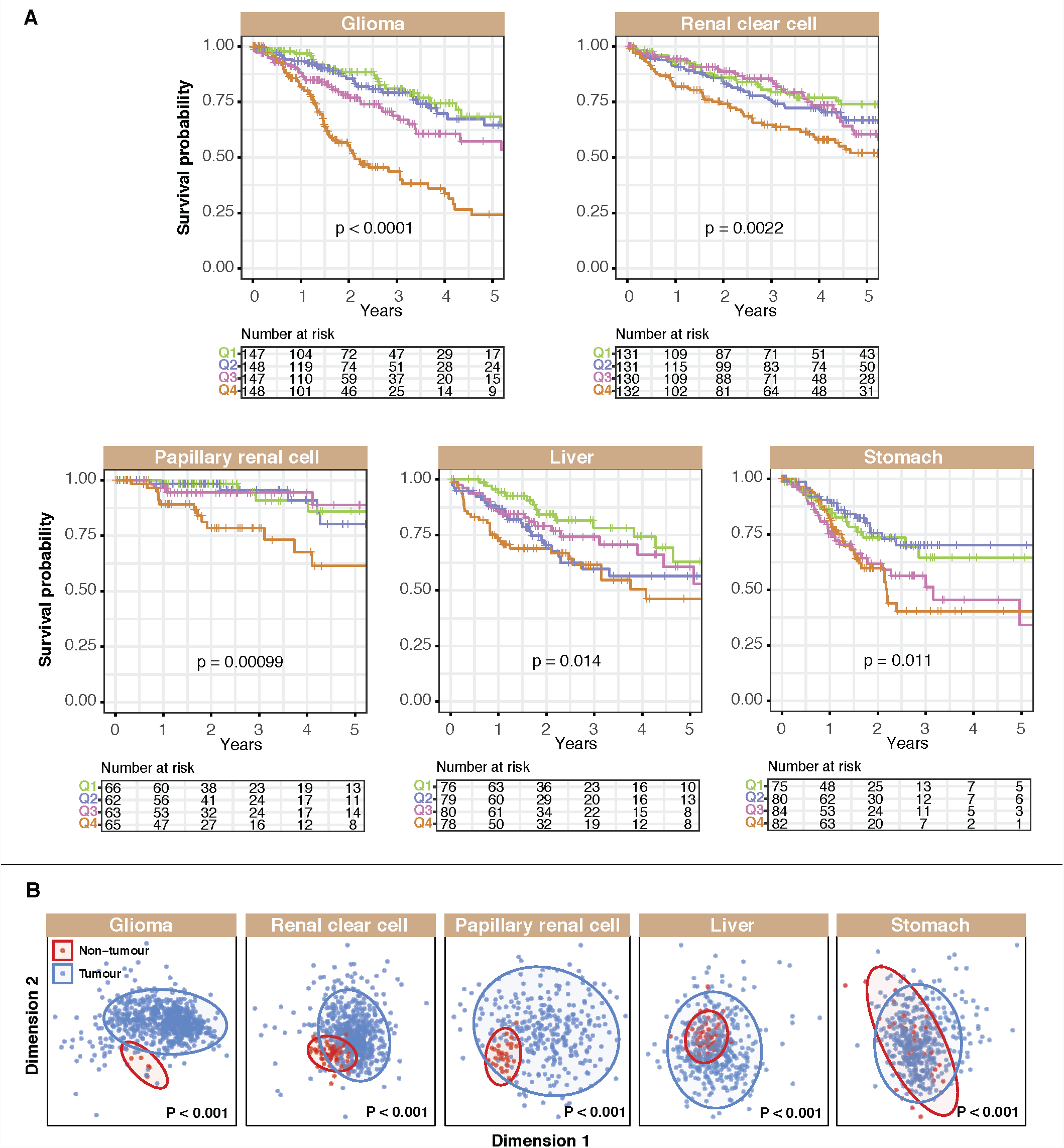
The Notch-Hedgehog 13-gene signature predicts patient survival in five cancers. **(A)** Kaplan-Meier estimates for overall survival using the signature. Patients were ranked and quartile stratified into Q1 (<25%), Q2 (25-50%), Q3 (50-75%) and Q4 (>75%) based on their 13-gene scores. P values were determined using the log-rank test. (**B)** Separation of tumour from non-tumour samples using the signature. Ordination plots of MDS analysis of the signature using Euclidean distances to represent tumour and non-tumour samples in 2-dimensional space. PERMANOVA test confirmed statistically significant differences between tumour and non-tumour samples.

Multivariate Cox proportional hazards regression was used to determine whether the signature was confounded by other clinicopathological features. Tumour, node, metastasis (TNM) staging is frequently used for patient stratification. Even after accounting for TNM staging, the signature remained an independent predictor of survival: clear cell renal cell (HR=1.731, P=0.014), papillary renal cell (HR=2.297, P=0.042), liver (HR=2.146; P=0.024) and stomach (HR=2.161, P=0.017) (Table S3). Given that both the signature and tumour stage were independent of each other, we reason that the signature could be used to improve TNM staging. We observed that Notch-Hedgehog driver genes offered an additional resolution in tumour classification for further stratification of similarly staged tumours in these cancers: clear cell renal cell (P<0.0001), papillary renal cell (P<0.0001), liver (P<0.0001) and stomach (P=0.0068) (Fig. 3A).

**Figure 3.**
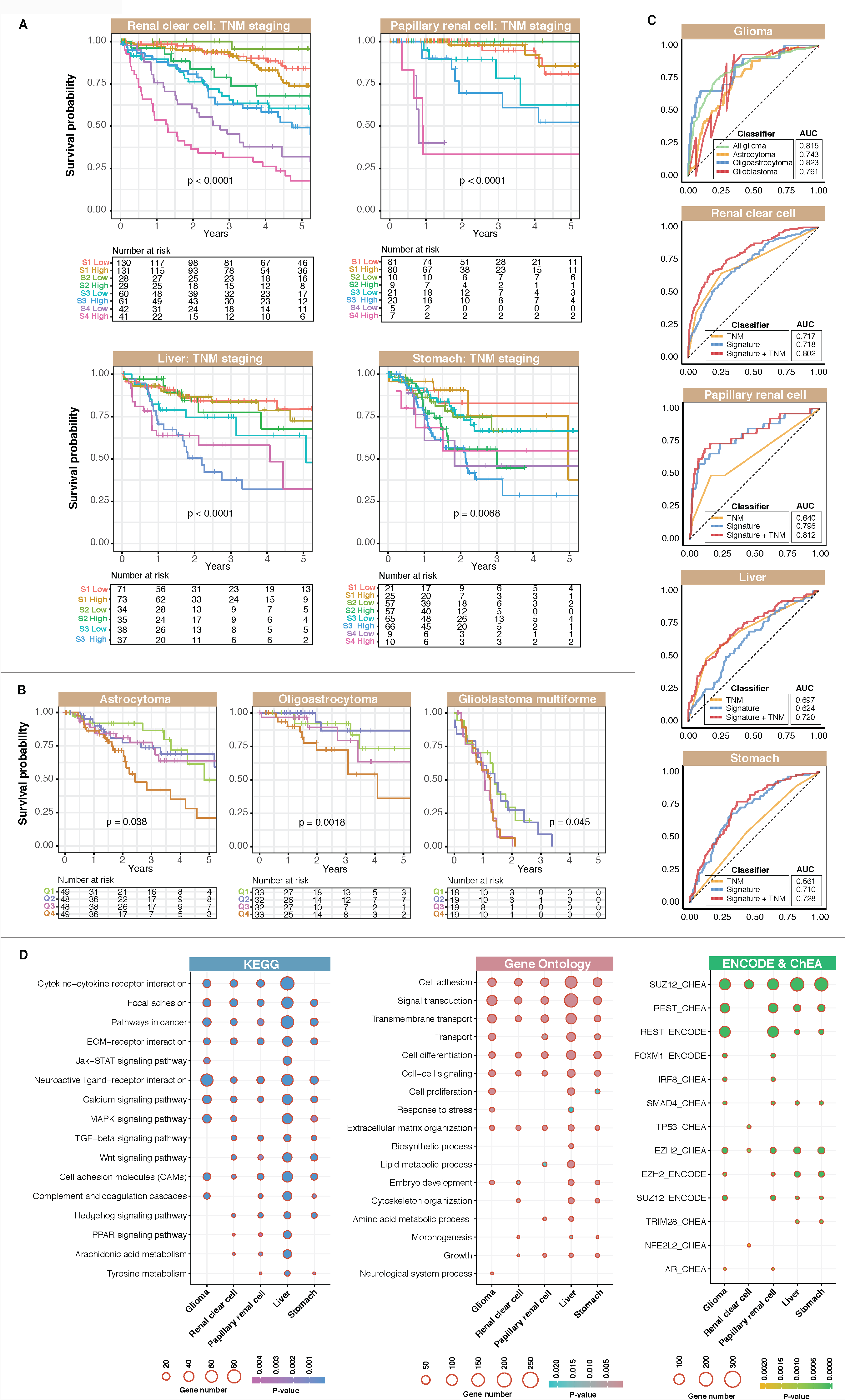
The Notch-Hedgehog 13-gene signature is independent of TNM stage and predicts overall survival in glioma histological subtypes. **(A)** Kaplan-Meier analyses were performed on patients stratified according to TNM stages and 13-gene scores. Patients were first separated into TNM stage and then median-stratified into low-and high-score groups based on their 13-gene scores. P values were determined using the log-rank test. (**B)** Kaplan-Meier estimates for overall survival using the signature on glioma subtypes ranging from low-grade (astrocytoma, oligoastrocytoma) to high-grade gliomas (glioblastoma multiforme). Patients were first stratified by histological subtypes followed by quartile stratification into Q1 (<25%), Q2 (25-50%), Q3 (50-75%) and Q4 (>75%) based on their 13-gene scores. P values were determined using the log-rank test. (**C)** Predictive performance of the signature. The receiver operating characteristic (ROC) analysis was used to assess specificity and sensitivity of the signature in predicting 5-year overall survival. ROC curves generated from the signature were compared to those generated from TNM staging and a combined model uniting TNM stage and the signature. AUCs for TNM stage were in accordance with previous publications employing TCGA datasets^1,22,47^. AUC: area under the curve. TNM: tumour, node and metastasis. (**D)** Enriched biological pathways and transcription factor binding associated with DEGs. Differential expression analyses were performed between Q4 and Q1 patients followed by mapping of DEGs against KEGG, Gene Ontology, ChEA and ENCODE databases.

Glioma samples are classified into four histological categories with varying severity: low-grade astrocytoma, low-grade oligodendroglioma, low-grade oligoastrocytoma (consisting of both abnormal astrocytoma and oligodendroglioma cells), and grade IV glioblastoma multiforme. Kaplan-Meier analyses of glioma samples grouped by histology revealed that the signature remained prognostic in astrocytoma (P=0.038), oligoastrocytoma (P=0.0018) and glioblastoma multiforme (P=0.045) (Fig. 3B). Patients with low-grade gliomas stratified by the signature into the 4^th^ quartile had significantly higher death risks compared to those within the 1^st^ quartile: astrocytoma (HR=2.535, P=0.021), oligoastrocytoma (HR=4.169, P=0.014) and glioblastoma multiforme (HR=2.163, P=0.042) (Table S3).

To evaluate the predictive performance of the signature, we employed receiver operating characteristic (ROC) analyses and compared area under the curves (AUCs) derived from the signature versus those derived from TNM staging. The signature had greater sensitivity and specificity in predicting 5-year overall survival compared to TNM staging: papillary renal cell (AUC=0.796 vs. AUC 0.640) and stomach (AUC=0.710 vs. AUC=0.561) (Fig. 3C). Importantly, when used as a combined model with TNM staging, it outperformed either the signature or TNM when considered alone, suggesting that incorporating molecular subtype information on Notch-Hedgehog signalling allowed more precise stratification: clear cell renal cell (AUC=0.802), papillary renal cell (AUC=0.812), liver (AUC=0.720) and stomach (AUC: 0.728) (Fig. 3C). In terms of predicting prognosis in glioma subtypes, performance of the signature was the best in oligoastrocytoma (AUC=0.823), followed by glioblastoma multiforme. (AUC=0.761) and astrocytoma (AUC=0.743) (Fig. 3C). The signature also performed well when all glioma subtypes were considered as a group (AUC=0.815) (Fig. 3C).

### The Notch-Hedgehog signature identifies molecular subtypes with stem cell-like features

Notch-Hedgehog hyperactivation is associated with increased mortality rates (Fig. 2, 3). To further investigate the underlying biological consequences of augmented Notch-Hedgehog signalling and how they lead to adverse outcomes, we performed differential expression analyses on all transcripts comparing high-and low-risk patients as predicted by the 13-gene signature. The liver cancer cohort had the highest number of differentially expressed genes (DEGs): 3,015 genes (-1.5 > log_2_ fold change > 1.5; P<0.01) (Table S4). This was followed by glioma (1,407 genes), stomach (906 genes), papillary renal cell (817 genes) and clear cell renal cell (545 genes) carcinoma (Table S4). Despite having very different pathologies, there was a great deal of DEG overlap between these cancers. 14 DEGs were found in all five cancers, 164 DEGs were observed in at least four cancers and 470 DEGs in at least three cancers (Fig. S3A), implying conserved biological roles of Notch-Hedgehog signalling in driving disease progression.

KEGG pathway analyses on DEGs demonstrated enrichments of pathways involved in regulating self-renewal and pluripotency, i.e. Wnt, TGF-β, MAPK, JAK-STAT and PPAR signalling (Fig. 3D; Fig. S3B), suggesting that tumours with hyperactive Notch-Hedgehog signalling were characterised by molecular footprints of stemness and that there was significant crosstalk between Notch-Hedgehog and other pathways involved in controlling tumour initiation. Additionally, Gene Ontology analyses revealed significant enrichments of processes related to cell differentiation, cell proliferation, embryo development and morphogenesis (Fig. 3D), supporting the hypothesis that tumour aggression and elevated mortality could be caused by the presence of cancer stem cells (CSCs) that are likely to be refractory to therapy. Consistent with these results, Enrichr transcription factor (TF) analyses revealed that TFs associated with stem cell function appeared amongst top enriched candidates (Fig. 3D). DEGs were enriched as binding targets of SUZ12, REST, EZH2, SMAD4 and FOXM1 as supported by both ChEA and ENCODE databases (Fig. 3D). Binding targets of SUZ12 and EZH2 were consistently enriched across all five cancer types, while targets of REST and SMAD4 were enriched in all cancers except for clear cell renal cell carcinoma (Fig. 3D). These TFs were thought to induce epithelial-mesenchymal transition and promote invasion and metastasis consistent with their roles in tumour initiation and maintenance^26^–^28^.

To independently confirm that the 13-gene signature is a potential pan-cancer marker of CSCs, we performed Spearman’s correlation analyses to compare 13-gene scores with expression profiles of other CSC markers where we would expect to see positive correlations. We examined expression profiles of nine genes implicated in CSC regulation: *CD105, CD133, CD200, CD24, CD29, CD44, CD73, CD90* and *NESTIN*. Putative neural CSC markers are *CD133, NESTIN, CD105* and *CD44*^29^. We observed significant positive correlations between 13-gene scores and all four markers in glioma samples (Fig. S4). *CD105, CD29, CD44, CD73, CD90* and *NESTIN* were positively correlated with 13-gene scores in renal cancers (Fig. S4); an observation which is consistent with these genes being markers of renal CSCs^30^. Seven and four CSC markers were positively correlated with 13-gene scores in liver and stomach cancers respectively (Fig. S4). Given the tissue-specific nature of these genes, we would not expect to see positive correlations in all cases. Nonetheless, our results overall suggest that hyperactive Notch-Hedgehog signalling is associated with CSC phenotypes, contributing to tumour aggression and poor survival outcomes.

### Transcription factors involved in self-renewal processes influence survival outcomes in patients with hyperactive Notch-Hedgehog signalling

Previously, we observed that binding targets of TFs associated with stem cell function were enriched amongst DEGs (Fig. 3D). Polycomb proteins, EZH2 and SUZ12 have been implicated in CSC formation and maintenance^31,32^. REST is a transcriptional repressor involved in maintaining embryonic and neural stem cell phenotypes^33^. Given their roles in CSC maintenance, we would expect to see elevated expression of these TFs in tumours with hyperactive Notch-Hedgehog signalling. Indeed, we observed significant positive correlations between 13-gene scores and *EZH2* levels in glioma (rho=0.45; P<0.0001), clear cell renal cell (rho=0.22; P<0.0001), papillary renal cell (rho=0.33; P<0.0001) and liver (rho=0.26; P<0.0001) cancers (Fig. 4A). Additionally, in the glioma cohort, positive associations between 13-gene scores and *REST (*rho=0.39; P<0.0001) or *SUZ12 (*rho=0.17; P<0.0001) profiles were observed (Fig. 4D).

**Figure 4.**
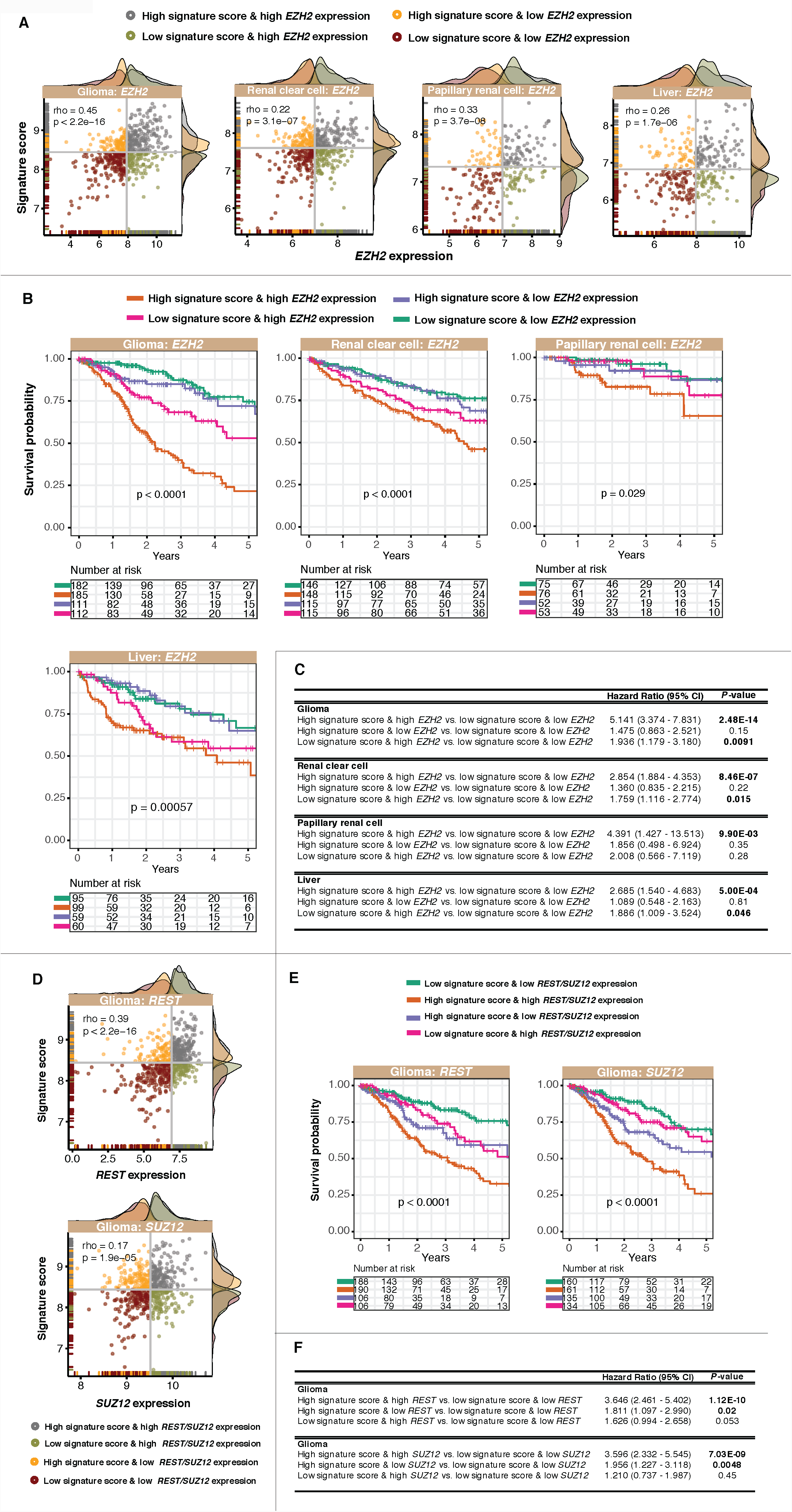
Prognostic significance of a combined model of the Notch-Hedgehog signature and transcription factors (*EZH2, SUZ12* and *REST*) involved in pluripotency maintenance. **(A)** Scatter plots demonstrate significant positive correlations between 13-gene scores and *EZH2* expression profiles in four cancers. Patients were stratified into four categories based on median 13-gene scores and *EZH2* expression. Density plots depict the distribution of 13-gene scores and *EZH2* expression at the y-and x-axes respectively. (**B)** Kaplan-Meier analyses were performed on the four patient categories to ascertain the combined relationship of the signature and *EZH2* expression on overall survival. (**C)** Table inset depicts univariate Cox proportional hazards analyses of the relationship between *EZH2* and the signature in four cancer types. (**D)** Scatter plots demonstrate significant positive correlations between 13-gene scores and *REST* or *SUZ12* expression levels in glioma. Patients were stratified into four categories based on median 13-gene score and *REST* or *SUZ12* expression. Density plots depict the distribution of 13-gene scores and *REST* or *SUZ12* expression at the y-and x-axes respectively. (**E)** Kaplan-Meier analyses were performed on the four patient categories to ascertain the combined relationship between the signature and *REST* or *SUZ12* expression on overall survival in glioma. (**F)** Table inset depicts univariate Cox proportional hazards analyses of the relationship between *REST* or *SUZ12* and the signature in glioma. CI: confidence interval. Significant P values are highlighted in bold.

To determine whether these associations harboured prognostic information, patients were categorised by their 13-gene scores and expression profiles of individual TFs into four categories: *1)* high 13-gene score and high TF expression, *2)* high 13-gene score and low TF expression, *3)* low 13-gene score and high TF expression and *4)* low 13-gene score and low TF expression (Fig. 4A and 4D). Interestingly, combined relationship of the signature and TF expression profiles allowed further delineation of patients into additional risk groups: glioma (*EZH2*: P<0.0001; *REST*: P<0.0001 and *SUZ12*: P<0.0001), clear cell renal cell (*EZH2*: P<0.0001), papillary renal cell (*EZH2*: P=0.029) and liver (*EZH2*: P<0.00057) cancers (Fig. 4B and 4E). Patients with high 13-gene scores that concurrently harboured high TF expression had the poorest survival outcomes: glioma (*EZH2*: HR=5.141, P<0.0001; *REST*: HR=3.646, P<0.0001; *SUZ12*: HR=3.596, P<0.0001), clear cell renal cell (*EZH2*: HR=2.854, P<0.0001), papillary renal cell (*EZH2*: HR=4.391, P=0.0099) and liver (*EZH2*: HR=2.685, P=0.0005) cancers (Fig. 4C and 4F). Taken together, our results suggest that coregulation by Notch-Hedgehog signalling and CSC TFs could synergistically contribute to more advanced disease states.

### Tumour hypoxia exacerbates disease phenotypes in Notch-Hedgehog^+^ CSCs

Hypoxia is intricately linked to pluripotency as it promotes stem cell maintenance and self-renewal in both embryonic stem cells and CSCs^34^, in part, through modulating hypoxia-inducible factor (HIF) function^35^. For example, glioma stem cells are typically found in the vicinity of necrotic regions that are hypoxic^36^. Glioma stem cells have increased ability to stimulate angiogenesis through VEGF upregulation^37^ and inhibition of HIFs could reduce CSC survival, self-renewal and proliferation^36^. We reason that hypoxia functions to maintain CSC niches. To assess the levels of tumour hypoxia, we employed a 52-hypoxia gene signature^17^ for calculating hypoxia scores in each patient by taking the average expression of hypoxia signature genes^17^. Indeed, we observed significant positive correlations between Notch-Hedgehog^+^ CSCs and hypoxia scores in glioma (rho=0.33, P<0.0001) and clear cell renal cell carcinoma (rho=0.16, P=0.00031) (Fig. 5A). By grouping patients based on their 13-gene and hypoxia scores, this joint model allowed the identification of patients with potentially more hypoxic tumours harbouring Notch-Hedgehog^+^ CSCs, which influenced overall survival rates: glioma (P<0.0001) and clear cell renal cell carcinoma (P=0.00013) (Fig. 5B). Indeed, patients with high CSC and hypoxia scores had significantly poorer survival outcomes: glioma (HR=6.008; P<0.0001) and clear cell renal cell carcinoma (HR=2.389, P<0.0001) (Fig. 5C). The CSC-hypoxia model is also prognostic in glioma subtypes: astrocytoma (HR=5.052, P<0.0001), oligoastrocytoma (HR=16.717, P=0.0066) and glioblastoma (HR=2.686, P=0.022) (Fig. 5B and 5C). Our results suggest that hypoxic zones within tumours could very well represent CSC niches.

**Figure 5.**
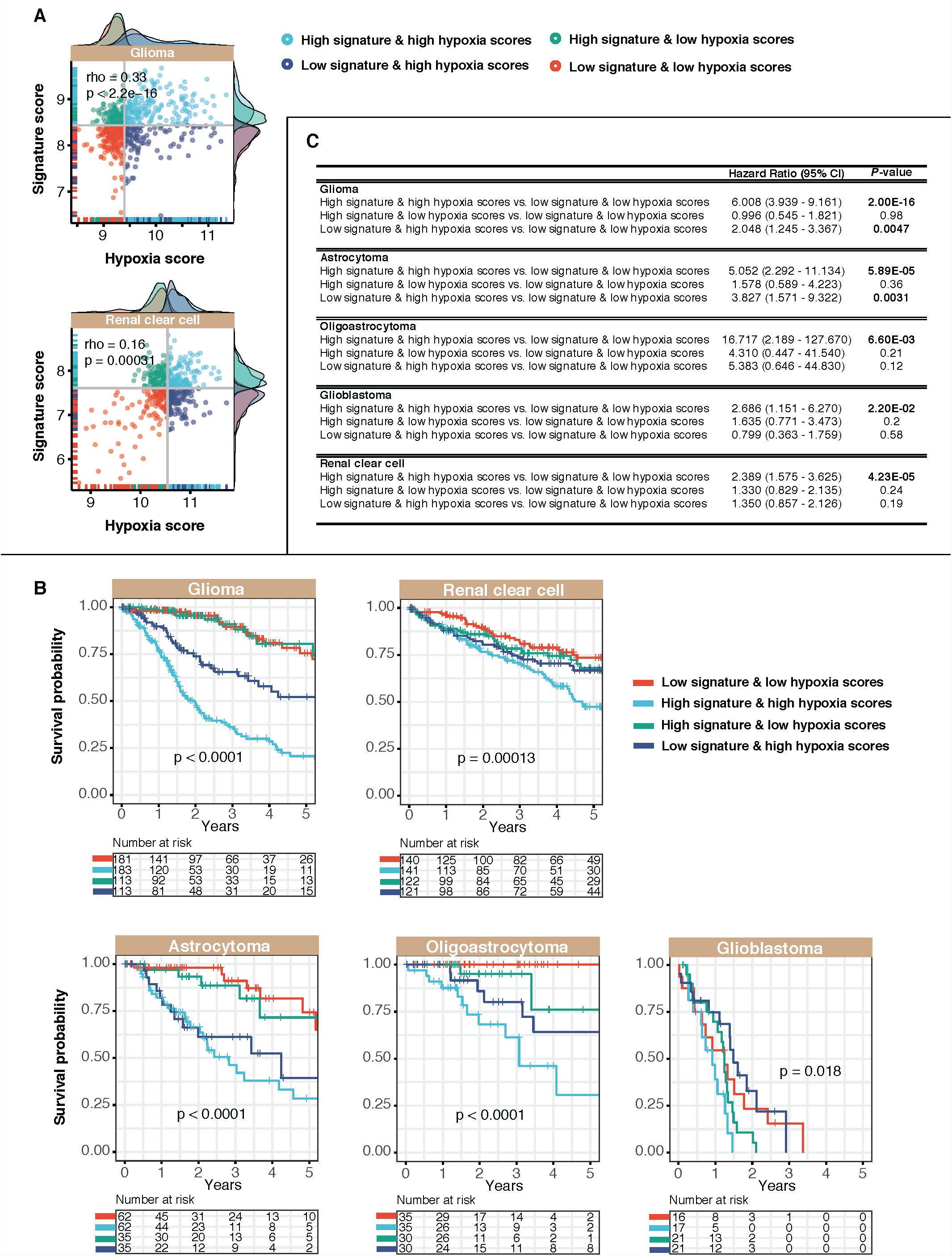
Positive associations between Notch-Hedgehog^+^ CSCs and tumour hypoxia in glioma and clear cell renal cell carcinoma. **(A)** Scatter plots demonstrate significant positive correlations between 13-gene and hypoxia scores. Patients were stratified into four categories based on median 13-gene and hypoxia scores. Density plots depict the distribution of 13-gene and hypoxia scores at the y-and x-axes respectively. (**B)** Kaplan-Meier analyses were performed on the four patient categories to ascertain the combined relationship of the signature and tumour hypoxia on overall survival. Contribution of hypoxia on Notch-Hedgehog^+^ CSCs were also determined in histological subtypes of glioma (astrocytoma, oligoastrocytoma and glioblastoma multiforme). (**C)** Table inset demonstrates univariate Cox proportional hazards analyses of the relationship between tumour hypoxia and the signature in glioma and clear cell renal cell carcinoma. CI: confidence interval. Significant P values are highlighted in bold.

### Putative Notch-Hedgehog^+^ CSCs are potentially immune privileged

Cancer progression is negatively correlated with immunocompetence of the host and evidence points to the role of CSCs in immunomodulation^3,38^. CSCs reside within niches that are often protected from environmental insults as well as attacks by the immune system. Hypoxic zones not only serve as CSC niches (Fig. 5)^39^, but also attract immunosuppressive cells such as regulatory T cells (Tregs)^22,40^, tumour-associated macrophages^41^ and myeloid-derived suppressor cells^42^. Given that positive associations between Notch-Hedgehog^+^ CSCs and hypoxia were linked to poor progression in glioma and clear cell renal cell carcinoma, we hypothesize that tumours characterised by these features would be immune privileged or hypoimmunogenic.

To test this hypothesis, we retrieved a list of 31 genes that represent tumour-infiltrating Tregs. This gene list was identified from the overlap of four Treg signatures^18^–^21^ to yield a more representative profile of tumour-infiltrating Tregs that is not specific to a single cancer type. A Treg score for each patient within the glioma and clear cell renal cell carcinoma cohorts was calculated as the mean expression of the 31 genes. We observed significant positive correlations between Treg scores and the Notch-Hedgehog 13-gene scores in both cohorts, supporting the hypothesis that CSCs are potentially hypoimmunogenic: glioma (rho=0.43; P<0.0001) and clear cell renal cell carcinoma (rho=0.31; P<0.0001) (Fig. 6A). As performed previously, patients were separated into four groups based on their 13-gene and Treg scores. When used in combination with the Notch-Hedgehog signature, Treg expression profiles allowed further separation of patients into additional risk groups that influenced overall survival: glioma (P<0.0001) and clear cell renal cell carcinoma (P<0.0001) (Fig. 6B). Intriguingly, patients characterised by high 13-gene and Treg scores had significantly higher mortality rates compared to those with low 13-gene and Treg scores: glioma (HR=4.921, P<0.0001) and clear cell renal cell carcinoma (HR=2.968, P<0.0001) (Fig. 6C). This was also true for other histological subtypes of glioma: astrocytoma (HR=2.721; P=0.0032), oligoastrocytoma (HR=5.431; P=0.0091) and glioblastoma (HR=3.065; P=0.0068) (Fig. 6C). Taken together, our results suggest that CSCs found within immunosuppressed environments are likely to be more aggressive.

**Figure 6.**
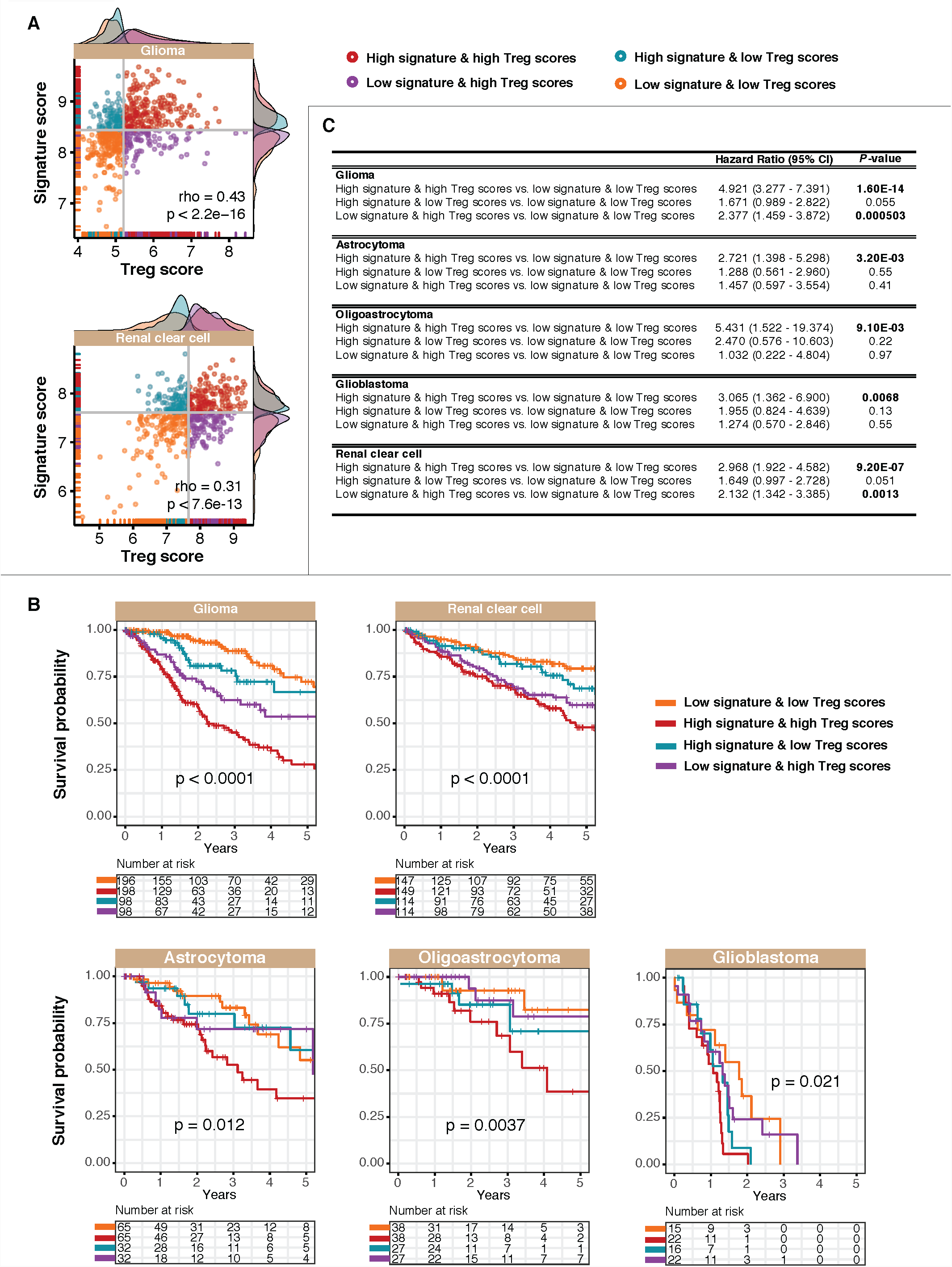
Positive associations between Notch-Hedgehog^+^ CSCs and immunosuppression in glioma and clear cell renal cell carcinoma. **(A)** Scatter plots demonstrate significant positive correlations between 13-gene and Treg scores. Patients were stratified into four categories based on median 13-gene and Treg scores. Density plots depict the distribution of 13-gene and Treg scores at the y-and x-axes respectively. (**B)** Kaplan-Meier analyses were performed on the four patient categories to ascertain the combined relationship of the signature and Treg-mediated immunosuppression on overall survival. Contribution of Tregs on Notch-Hedgehog^+^ CSCs were also determined in histological subtypes of glioma (astrocytoma, oligoastrocytoma and glioblastoma multiforme). (**C)** Table inset demonstrates univariate Cox proportional hazards analyses of the relationship between Tregs and the signature in glioma and clear cell renal cell carcinoma. CI: confidence interval. Significant P values are highlighted in bold.

## Discussion and Conclusion

Aberrations in the Notch-Hedgehog signalling axis are frequently implicated in malignant progression. Hedgehog genes, *Shh, PTCH1* and *GLI1*, were detected in over 50% of liver cancer tumours and inhibition of Hedgehog signalling by cyclopamine, Smoothened antagonist or anti-SHH resulted in decreased cell growth and increased apoptosis^43^. Notch signalling is also activated in liver cancer and this leads to formation of liver tumours in mice^44^. Notch blockade using γ-secretase inhibitors reduced cell viability in hepatoma cell lines^44^. In clear cell renal cell carcinoma, inhibition of Notch signalling reduced anchorage-independent growth and mice treated with Notch inhibitors had impaired growth of transplanted cancer cells^45^. Elevated expression of Notch ligands correlated with aggressiveness and poor survival rates in stomach cancer^46^.

These studies have paved the road for understanding the role of Notch-Hedgehog signalling in carcinogenesis. However, large-scale comparative studies investigating the similarities and differences in Notch-Hedgehog signalling across multiple cancer types have remained limited. We interrogated expression and mutational profiles of 72 genes from Notch and Hedgehog pathways in 21 diverse cancer types involving 18,484 patients. Our integrated analysis of genomic, transcriptomic and clinical data revealed molecular distinct tumour subtypes that were characterised by Notch-Hedgehog hyperactivation. Concentrating on 13 Notch-Hedgehog driver genes that were recurrently amplified and overexpressed, we found that these genes were associated with clinically relevant molecular features of stemness. The biological consequences of elevated expression of driver genes were manifold. High-risk patients showed overexpression of genes associated with other stem cell-related pathways such as Wnt, JAK-STAT and TGF-β signalling (Fig. 3D. and S3B)^47^. Simultaneous inhibition of Notch and JAK-STAT pathways by combined AG-490 and GSI IX therapy impaired pancreatic cancer progression^48^. *GLI2* is regulated by both Hedgehog and TGF-β pathways and others have surmised that TGF-β may potentiate Hedgehog signalling cascade by increasing *GL12* availability, contributing to metastasis^49^. Hence, our study reveals molecular targets with overlapping functions that can be prioritised to improve therapeutic outcomes.

Furthermore, binding targets of stem cell-related TFs (*EZH2, SUZ12* and *REST*) were enriched amongst genes upregulated in high-risk patients (Fig. 4). *EZH2* synergises with Notch-Hedgehog^+^ CSCs to worsen survival outcomes in patients with glioma, renal and liver cancers (Fig. 4). Pharmacological inhibition of *EZH2* impaired glioblastoma CSC tumour-initiating capacity and survival^31^. *EZH2*-mediated transcriptional silencing leads to the maintenance of undifferentiated states in glioblastoma through *STAT3* activation^50^. In liver cancer, *EZH2* overexpression is associated with vascular invasion, malignant progression^51^ and activation of β-catenin/Wnt signalling^52^. Inhibition of *EZH2* in renal cancer cell lines led to increased apoptosis^53^. Additionally, enrichments of *SUZ12* and *REST* targets in glioma patients with hyperactive Notch-Hedgehog signalling were linked to significantly poorer prognosis (Fig. 4D and 4E). *REST* is implicated in transcriptional regulation of neuronal stem cells^33^, while the overexpression of *SUZ12* is linked to tumour progression^54^.

An exploration of the relationship between Notch-Hedgehog hyperactivation and tumour microenvironmental qualities revealed associations of CSCs with hypoxia and immunosuppression. We observed that CSCs characterised by hyperactive Notch-Hedgehog signalling exhibited immune privileged features associated with the attenuation effects of Tregs (Fig. 6). Effectiveness of immunotherapy is biased towards differentiated cells that make up the tumour bulk due to distinct antigen presentation in CSCs^55^. CD133^+^ glioma CSCs fail to express NK cell ligands or MHCI, which prevents immune detection^56^. Stimulatory NK cell ligands are also downregulated in breast CSCs, contributing to evasion from NK cell killing^57^. Pan et al. elegantly reviewed recent initiatives focusing on immunotherapeutic agents against CSC antigens employing dendritic cell vaccines, myeloid-derived suppressor cell-based approaches and the use of immune checkpoint blockades recognising *PD-1* or *CTLA4*^55^. The Notch-Hedgehog signature may be used to stratify patients prior to immunotherapy.

Immunoevasion can be exacerbated by tumour hypoxia as the latter not only promotes CSC survival, but also creates an environment that facilitates further immune suppression^22^. It may be possible that Notch-Hedgehog^+^ CSCs in glioma and clear cell renal cell carcinomas are more frequently found within immunosuppressed hypoxic zones (Fig. 5). Indeed, hypoxia could stimulate self-renewal of CD133^+^ glioma stem cells and this is abrogated by HIF-1α knockdown^58^. Hypoxia promotes the maintenance of undifferentiated states through the activation of Notch-responsive genes in neuronal progenitors^35^. Hypoxia also activates cellular reprogramming of non-stem cancer cells into CSCs in glioblastoma by inducing the expression of *Oct4, Nanog* and *c-Myc*^59^. Glioma stem cells are pro-angiogenic due to promiscuous secretions of VEGF that is further induced by hypoxia^37^. Bevacizumab, which targets VEGF, could suppress xenographs derived from glioma stem cells but not those derived from non-stem glioma cells^37^. In renal cancer, we observed that Notch-Hedgehog^+^ CSCs are likely to be enriched in hypoxic tumours and the combined effects of hypoxia and augmented Notch-Hedgehog signalling resulted in further elevation of death risks (Fig. 5). However, Sjölund et al. observed that Notch signalling is not enhanced by hypoxia in renal cancer^45^. Another study on renal CSCs revealed that hypoxia did not affect the differentiation potential of CD105^+^ CSCs^60^. Nonetheless, hypoxia was found to induce the expression of stem cell markers, *Oct4, Nanog, c-Myc* and *Klf4* in renal cancer cell lines, supporting our observation, and in another ten cancers including cervix, lung, colon, liver and prostate^61^.

Although prospective validation is warranted, the results presented in this work support a model where Notch-Hedgehog hyperactivation is linked to stemness and that hypoxia contributes to the maintenance of undifferentiated phenotypes and the reduction of anti-tumour immunity. The use of immune checkpoint blockade has been increasingly tried in malignancy^62^. Hence, molecular signatures capable of discerning responders from non-responders will be valuable prior to the administration of these expensive drugs. As an independent prognostic indicator in five cancer types involving 2,278 patients, the Notch-Hedgehog gene signature may serve as a staging point for exploring combinatorial treatments that simultaneously target CSCs, hypoxia and tumour immunity.

## Additional Information

### Ethics approval and consent to participate

Not applicable.

### Consent for publication

Not applicable.

### Availability of data and material

The datasets supporting the conclusions of this article are included within the article and its supplementary files.

### Conflict of interest

None declared.

### Funding

None.

### Authors’ contributions

AGL designed the study and supervised the research. WHC and AGL analysed the data, interpreted the results and wrote the initial manuscript draft. AGL revised the manuscript draft. All authors approved the final submitted manuscript.

## Supplementary figure and table legends

**Figure S1. Prognosis of each of the 13 signature genes in 20 cancer types determined using Cox regression analyses.** Rows (Notch-Hedgehog driver genes) and columns (cancer types) were ordered using hierarchical clustering (Euclidean distance metric). Boxes in the lightest pink colour represent non-prognostic genes. Heatmap intensities represent hazard ratios of prognostic genes that were significant (P<0.05).

**Figure S2. Expression distribution of the 13 signature genes in tumour and non-tumour samples of five cancers.** Nonparametric Mann-Whitney-Wilcoxon tests were employed to determine whether there were significant differences in expression distributions. Asterisks represent significant P values: * < 0.05, ** < 0.005, ***<0.0005 and ****<0.00005. ns: non-significant.

**Figure S3. Differentially expressed genes between Q4 and Q1 patients as determined by their 13-gene scores. (A)** Venn diagram depicts a five-way comparison of DEGs (-1.5 > log_2_ fold-change > 1.5, P<0.01) identified from five cancer cohorts. (**B)** Volcano plots illustrate the distribution of DEGs (in pink). Non-significant genes were represented as grey dots. Genes implicated in other stem cell-related signalling modules were annotated and colour-coded.

**Figure S4. Correlations between the Notch-Hedgehog signature and other CSC markers.** Scatter plots depict the associations between 13-gene scores and nine CSC marker expression profiles in five cancer types. P values were determined by Spearman’s rank-order correlation analyses.

**Table S1.** List of 72 Notch-Hedgehog pathway genes.

**Table S2.** List of the number of tumour and non-tumour samples obtained from TCGA along with abbreviation decodes.

**Table S3.** Univariate and multivariate Cox proportional hazards regression to determine the independence of the signature with other clinicopathological risk factors. Significant P values were highlighted in bold.

**Table S4.** Differentially expressed genes between Q4 and Q1 patient groups as determined by the signature in five cancer types.

